# An elevated level of the mRNA exporter Mex67-Mtr2 in nuclear mRNPs impairs nuclear mRNA export

**DOI:** 10.1101/2025.02.26.640412

**Authors:** Nataliia Stefanyshena, Katja Sträßer

## Abstract

In eukaryotes, a crucial step in gene expression is the export of the mRNA out of nucleus. The conserved mRNA exporter Mex67-Mtr2 (NXF1-NXT1 in human) is a key factor in this process. In *S. cerevisiae*, it is recruited to the mRNA by the four export adaptors Hpr1, Nab2, Yra1 and Npl3 that play important yet poorly defined roles in nuclear mRNA export. Here, we show that an excess of Mex67 in nuclear mRNPs impairs nuclear mRNA export. *Δhpr1* cells have increased Nab2, Yra1 and Mex67 levels in nuclear mRNPs in addition to the export defect. Notably, overexpression of Nab2 or Yra1 in *Δhpr1* cells suppresses this export defect and reduces the Mex67 level in nuclear mRNPs to a WT level. Importantly, the increase in the Mex67 level in *Δhpr1* cells is not a consequence and thus the cause of the nuclear mRNA export defect. Therefore, precise regulation of the Mex67-Mtr2 level in nuclear mRNPs is essential for efficient nuclear mRNA export.

## Introduction

Gene expression is an essential process for every cell and organism. In eukaryotic cells, RNA polymerase II (RNAPII) first transcribes a protein-coding gene into an mRNA. Largely already co-transcriptionally the mRNA is processed by addition of a cap structure to its 5’ end, splicing of the introns and addition of a poly(A) tail after cleavage at its 3’ end^1,2^. Importantly, the mRNA is also packaged into a messenger ribonucleoprotein particle (mRNP) by nuclear RNA-binding proteins (RBPs), so-called mRNP components^1-3^. These processes as well as the involved proteins are well-conserved^3,4^. In the model organism *S. cerevisiae*, most likely all nuclear mRNP components are known^5^. Most of these proteins have multiple functions along the gene expression pathway.

The TREX complex couples transcription elongation to nuclear mRNA export, most likely by binding co-transcriptionally to the nascent mRNA and facilitating mRNP assembly^4,6^. It consists of the pentameric THO complex, which in turn is composed of Tho2, Hpr1, Mft1, Thp2 and Tex1, the SR-like proteins Gbp2 and Hrb1, and the mRNA export factors Sub2 and Yra1^6^. The cap binding complex (CBC), consisting of Cbc1 and Cbc2, binds to the 5’ end of the mRNA and promotes nuclear mRNA export by recruiting Yra1 and Npl3 to chromatin in a THO- and Sub2-independent manner^7^. Tho1 is a conserved nuclear mRNP component proposed to function complementarily to Sub2, as overexpression of either suppresses *Δhpr1* defects^8^. It is recruited to transcribed genes in a THO- and RNA-dependent manner^8^. CIP29, its homolog in humans, interact withs TREX and UAP56/DDX39, the human homolog of Sub2, and mutation of MOS11, the Tho1 homolog in plants, causes nuclear poly(A) RNA accumulation^9,10^. Thus, Tho1/CIP29/MOS11 is critical for mRNP assembly and nuclear mRNA export, though its exact mechanisms remain unclear. Several SR- and SR-like mRNP components are involved in packaging mRNAs into mRNPs^11^. In *S. cerevisiae*, the SR-like protein Npl3 plays roles in transcription, splicing, 3’ end processing and mRNA export^12-14^. The <u>n</u>uclear poly(<u>A</u>)-<u>b</u>inding protein Nab2 is an SR-like protein, which functions in poly(A) tail length control, nuclear mRNP assembly and export^15^. It is also recruited to the site of transcription by binding to the RNA^15^. Similarly, in human cells, several nuclear mRNP components are SR- and SR-like proteins and important for nuclear mRNA export^11^. The THSC complex (TREX2), consisting of Thp1, Sac3, Sus1, Cdc31 and Sem1, is also essential for nuclear mRNA export^16^. It interacts with NXF1-NXT1/Mex67-Mtr2 and the nuclear pore complex (NPC)^16^. However, the exact functions and mechanisms of THSC remain unclear. Finally, the mRNP is exported out of the nucleus by the conserved mRNA exporter Mex67-Mtr2 in *S. cerevisiae* / NXF1-NXT1 in human cells. Thus, Mex67-Mtr2/NXF1-NXT1 is essential for nuclear mRNA export^17^. As the intrinsic affinity of Mex67-Mtr2 for RNA is very low, four export adaptor proteins, Hpr1, Nab2, Yra1 and Npl3 recruit the mRNA exporter to the mRNA in *S. cerevisiae*^18-21^. In mammalian cells, TREX (via its components ALYREF and THOC5), CHTOP, UIF, several SR-proteins and ZC3H3 recruit NXF1-NXT1 to the mRNA^22^. Mex67-Mtr2/NXF1-NXT1 also binds to nuclear pore proteins and exports the mRNP out of the nucleus. On the cytoplasmic side of the NPCs, several mRNP components including Mex67-Mtr2 are dissociated from the mRNA rendering nuclear mRNA export unidirectional^23,24^.

Nuclear mRNP assembly and export can be regulated, e.g. under stress conditions^25-27^. Despite its importance, it is not known how an mRNA and the nuclear mRNP components assemble into an mRNP and what the precise functions of each mRNP component are. Specifically, it is not known why at least four different adaptor proteins exist in *S. cerevisiae*, how they work together and how they compensate for each other.

Here, we show that in addition to a low level also an increased level of the mRNA exporter Mex67-Mtr2 in nuclear mRNPs causes a nuclear mRNA export defect. Deletion of *HPR1*, coding for one of the adaptor proteins, causes a nuclear mRNA export defect and higher levels of Nab2 and Yra1, two of the other adaptors, and of Mex67 in nuclear mRNPs. Interestingly, overexpression of Nab2 or Yra1 in *Δhpr1* cells suppresses the mRNA export defect. Concomitantly, the Nab2 level in nuclear mRNPs increases, while the Mex67 level decreases to wild-type amounts. Importantly, we show that the increased Mex67 level in nuclear mRNPs in *Δhpr1* cells is the cause, not the consequence, of the nuclear mRNA export defect. Thus, the maintenance of an optimal Mex67 level in nuclear mRNPs is required for efficient nuclear mRNA export.

## Results

### Loss of TREX causes a nuclear mRNA export defect and higher levels of the mRNP components Nab2 and Yra1 and the mRNA exporter Mex67 in nuclear mRNPs

For nuclear mRNA export, the conserved mRNA exporter Mex67-Mtr2 / NXF1-NXT1 is recruited to the nuclear mRNA by adaptor proteins^11,17,28^. In *S. cerevisiae*, four adaptors exist: Hpr1, Nab2, Yra1 and Npl3^18-21^. The existence of several adaptor proteins as well as experimental evidence suggest different export pathways for different classes of mRNAs. However, the molecular consequences of the lack of one of these potentially redundant export adaptors is largely unknown, especially in terms of nuclear mRNP biogenesis. Thus, we assessed the effects of lack of Hpr1, one of these four adaptors. To do so, the non-essential gene coding for Hpr1 was deleted or Hpr1 depleted using the auxin-inducible degron system (Supplementary Fig. 1a, b). Loss of Hpr1 by deletion or depletion causes a growth defect (Supplementary Fig. 1c and ^29^) and a concomitant nuclear mRNA export defect (Supplementary Fig. 1d and ^6,30^). Interestingly, in the absence of Hpr1, Thp2 still interacts with Mft1, but not any more with Tho2 or Hpr1 (Supplementary Fig. 1e). This is consistent with a recently published cryo-EM-based structural model of the *S*.*c*. THO complex, in which Thp2 and Mft1 are intertwined in a coiled coil, but Tho2 and Tex1 have much smaller interfaces with Mft1-Thp2 and rather interact with each other and with Hpr1 (Supplementary Fig. 1f and ^31^). Thus, lack of Hpr1 leads to disassembly of the THO complex, and the effects observed for cells lacking Hpr1 are probably due to the absence of a functional THO complex.

To assess potential changes in nuclear mRNP composition that might result from loss of Hpr1 and, thus, a functional THO complex, we purified nuclear mRNPs by native pulldown of endogenous TAP-tagged Cbc2, the small subunit of CBC. Interestingly, the absence of Hpr1 causes higher total levels of Nab2 and Yra1, two of the three other export adaptors, and concomitantly higher levels of both proteins in nuclear mRNPs (Fig. 1a, b). This is consistent with the finding that the amount of Nab2 in nuclear mRNPs is already increased after depletion of Hpr1 for 1 h (Supplementary Fig. 1g, h). In contrast, the level of Npl3, the fourth adaptor, is neither increased in the *HPR1* deletion strain nor after Hpr1 depletion (Fig. 1a, b, Supplementary Fig. 1g, h). However, much to our surprise, the level of the mRNA exporter Mex67 increases in nuclear mRNPs in *Δhpr1* cells compared to wild-type (WT) cells (Fig. 1a, b), despite the fact that *Δhpr1* cells have an mRNA export defect.

**Figure 1.**
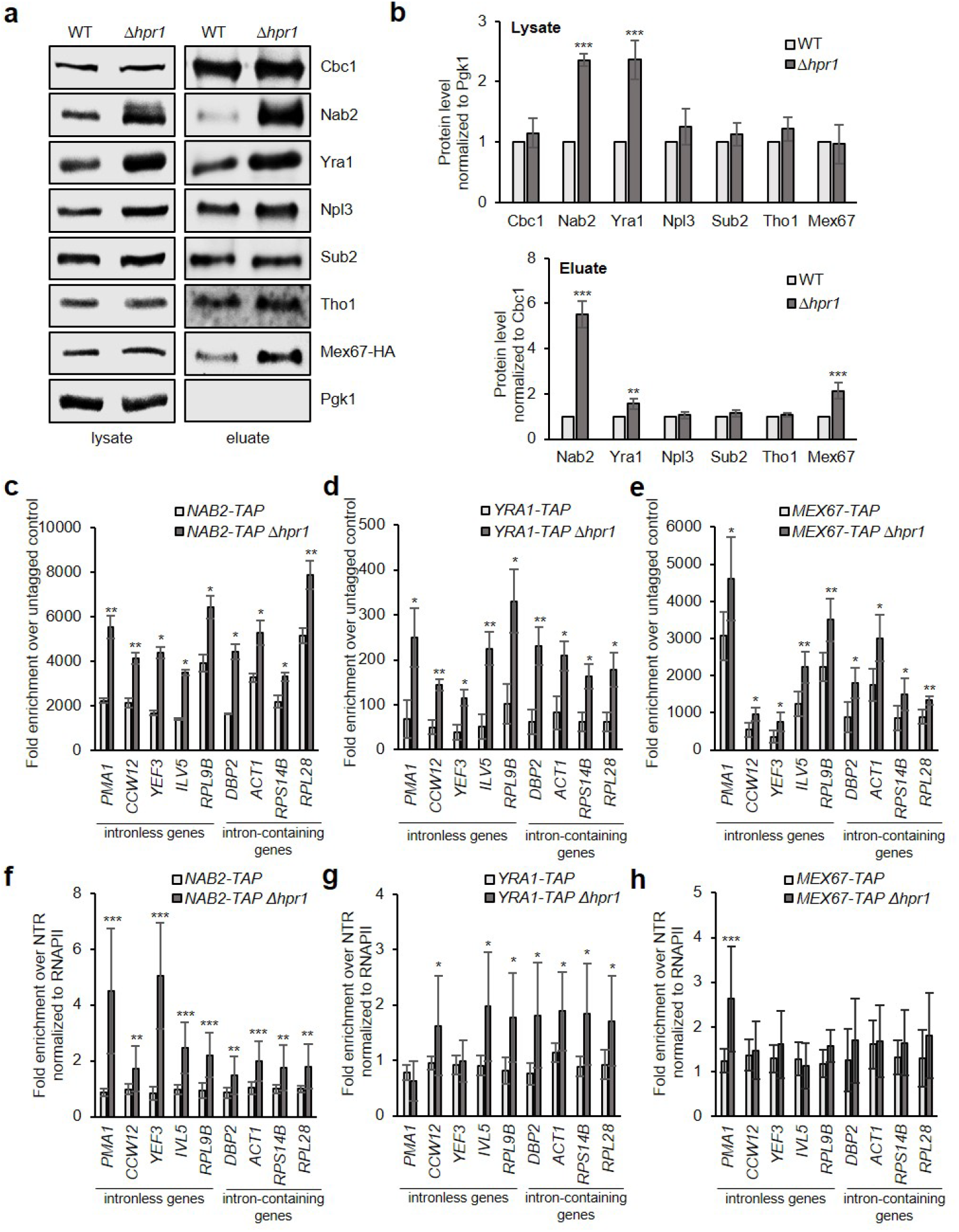
Loss of Hpr1 leads to increased Nab2, Yra1 and Mex67 levels in nuclear mRNPs. **a and b** The levels of Nab2, Yra1 and Mex67 in nuclear mRNPs are higher in *Δhpr1* cells. **a** Western blots of lysates and TEV eluates after Cbc2-TAP purification from WT and *Δhpr1* cells, using antibodies against the indicated RBPs. **b** Quantification of the protein levels in lysates (upper panel) and TEV eluates (lower panel), normalized to Pgk1 or Cbc1 levels, respectively. Values for WT cells were set to 1. **c - e** RNA binding of Nab2 (c), Yra1 (d) and Mex67 (e) increases in *Δhpr1* cells. TAP-tagged Nab2, Yra1 or Mex67 were immunoprecipitated and the amount of co-immunoprecipitated RNA was determined by RT-qPCR for representative abundant intronless and intron-containing transcripts. mRNA levels were quantified as the enrichment over the mRNA amounts in cells expressing untagged proteins. **f - h** The occupancy of Nab2 (f) and Yra1 (g) at transcribed genes is increased in *Δhpr1* cells, while the occupancy of Mex67 (h) remains unchanged. Chromatin immunoprecipitation (ChIP) analysis was used to assess the occupancy of TAP-tagged proteins at genes. Co-immunoprecipitated DNA was quantified by qPCR for the same intronless and intron-containing genes as in (**c - e**). The occupancy was calculated as the enrichment of protein at the genes relative to its presence at a non-transcribed region (NTR, 174131 - 174200 on chr. V), normalized to RNAPII occupancy and set to 1 for WT cells. Data represents the mean ± SD of at least three independent experiments; *p<0.05; **p<0.01; *** p<0.001.

The higher levels of Nab2, Yra1 and Mex67 in nuclear mRNPs in *Δhpr1* cells suggest that these proteins also bind more mRNA. To assess this, we performed RNA immunoprecipitation (RIP) experiments by purification of genomically TAP-tagged proteins from whole cell extracts from WT and *Δhpr1* cells (Fig. 1c-e). Coimmunoprecipitated RNA was analyzed by reverse transcription (RT) and quantitative PCR (qPCR) for five representative abundant intronless transcripts (*PMA1, CCW12, YEF3, ILV5* and *RPL9B*) and four intron-containing transcripts (*DBP2, ACT1, RPS14B* and *RPL28*). Indeed, the three proteins bind higher amounts of these mRNAs in the *Δhpr1* compared to the WT strain (Fig. 1c-e). Thus, nuclear mRNPs contain more Nab2, Yra1 and Mex67 in *Δhpr1* cells.

Nuclear mRNP components are recruited already to the transcribed gene. To determine whether the presence of Nab2, Yra1 and Mex67 is increased at genes in *Δhpr1* cells, we assessed their occupancy by chromatin immunoprecipitation (ChIP) (Fig. 1f-h). As the occupancy of nuclear RBPs depends on the presence of RNA and thus on transcription, we first assessed the occupancy of RNAPII at genes encoding the same nine representative transcripts assayed by RIP. RNAPII occupancy decreases in the *Δhpr1* strain (Supplementary Fig. 2). Accordingly, the occupancy of all RBPs was normalized to the one of RNAPII. Interestingly, the occupancy of Nab2 and Yra1, but not of the mRNA exporter Mex67 increases in *Δhpr1* cells (Fig. 1f-h). Taken together, lack of the THO complex leads to a higher occupancy of the adaptor proteins Nab2 and Yra1 at transcribed genes and to higher levels of these proteins in nuclear mRNPs. Potentially, these two adaptor proteins then increase the Mex67 level in nuclear mRNPs.

### Overexpression of Nab2 or Yra1 suppresses the growth and the nuclear mRNA export defect of *Δhpr1* cells

As a lack of Hpr1 leads to higher levels of Nab2, Yra1 and Mex67 in nuclear mRNPs, we hypothesized that overexpression of these proteins alleviates the growth and the nuclear mRNA export defect of *Δhpr1* cells. Overexpression of Npl3, whose level is not increased in nuclear mRNPs of *Δhpr1* cells, served as a potential negative control. As expected, overexpression of Npl3 in *Δhpr1* cells neither suppresses the growth nor the nuclear mRNA export defect of *Δhpr1* cells (Fig. 2a, b). Interestingly, overexpression of Nab2 or Yra1 suppresses the growth defect as well as the nuclear mRNA export defect of *Δhpr1* cells (Fig. 2a, b). Interestingly, overexpression of Mex67 does not suppress either defect (Fig. 2a, b). This is surprising, as the Mex67 level in nuclear mRNPs increases in *Δhpr1* cells and as Mex67-Mtr2 is the mRNA exporter. Taken together, overexpression of one of the two mRNA export adaptors Nab2 or Yra1 suppresses both defects indicating that the growth defect of *Δhpr1* cells is at least partially caused by their nuclear mRNA export defect.

**Figure 2.**
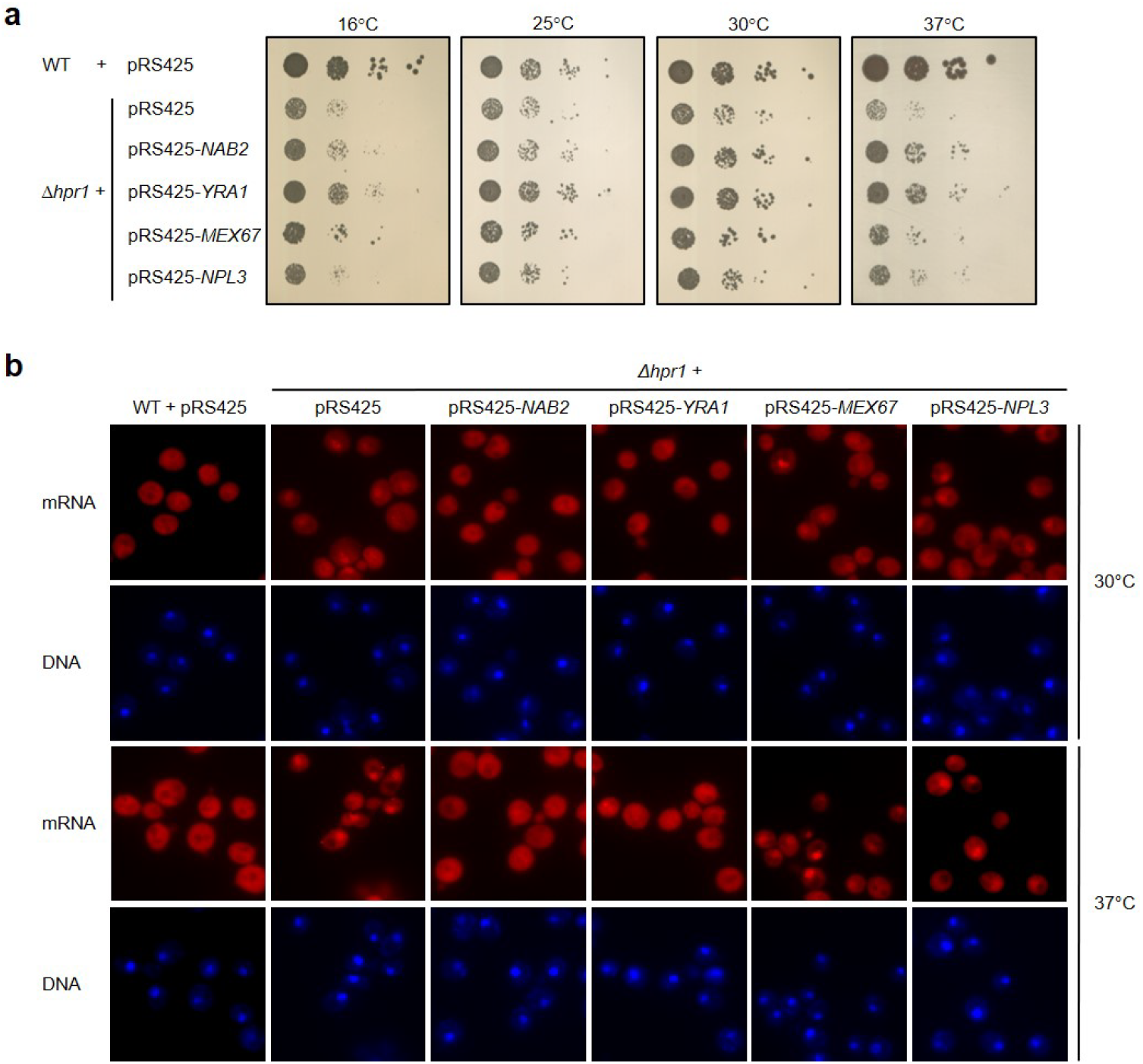
Overexpression of Nab2 or Yra1 suppresses the growth and nuclear mRNA export defects of *Δhpr1* cells. **a** The growth defect of *Δhpr1* cells is alleviated by overexpression of Nab2 or Yra1, but not by overexpression of Mex67 or Npl3. Ten-fold serial dilutions of WT or *Δhpr1* cells or *Δhpr1* cells overexpressing Nab2, Yra1, Mex67 or Npl3 were spotted onto SDS (−leu) plates and grown at the indicated temperatures. **b** The nuclear mRNA export defect of *Δhpr1* cells is suppressed by overexpression of Nab2 or Yra1, but not by overexpression of Mex67 or Npl3. Nuclear mRNA export was assessed by fluorescence in situ hybridization (FISH) of the indicated cells grown at 30°C or after a 1 h shift to 37°C. mRNA was detected using an oligo(dT)50 probe conjugated to the fluorescent dye Cy3. DNA was stained with DAPI. The results are representative for three independent experiments.

### Overexpression of Nab2 or Yra1 in *Δhpr1* cells increases Nab2 levels even further and decreases Mex67 levels in nuclear mRNPs

To the molecular basis for the suppression of the nuclear mRNA export defect in *Δhpr1* cells by overexpression of Nab2 or Yra1 we determined the composition of nuclear mRNPs of these cells (Fig. 3a, b). The total protein amount of Nab2 and Yra1 in *Δhpr1* cells increases further by overexpression of either Nab2 or Yra1 (Fig. 3a, b). Thus, the total amounts of these two proteins in cells are interdependent. Importantly, overexpression of Nab2 or Yra1 in *Δhpr1* cells further increases the already elevated Nab2 level in nuclear mRNPs, but not those of Yra1 (Fig. 3a, b). Surprisingly, the Mex67 level in nuclear mRNPs decreases to WT levels in *Δhpr1* cells, when either Nab2 or Yra1 are overexpressed (Fig. 3a, b). In contrast, the level of all assessed RBPs remains unchanged when Mex67 or Npl3 are overexpressed (Fig. 3a, b). Thus, when the nuclear mRNA export defect caused by the absence of Hpr1 is suppressed, the Nab2 level in nuclear mRNPs is even higher than in *Δhpr1* cells, and the Mex67 level decreases to WT. This suggests that a wild-type level of Mex67 is necessary for efficient nuclear mRNA export.

**Figure 3.**
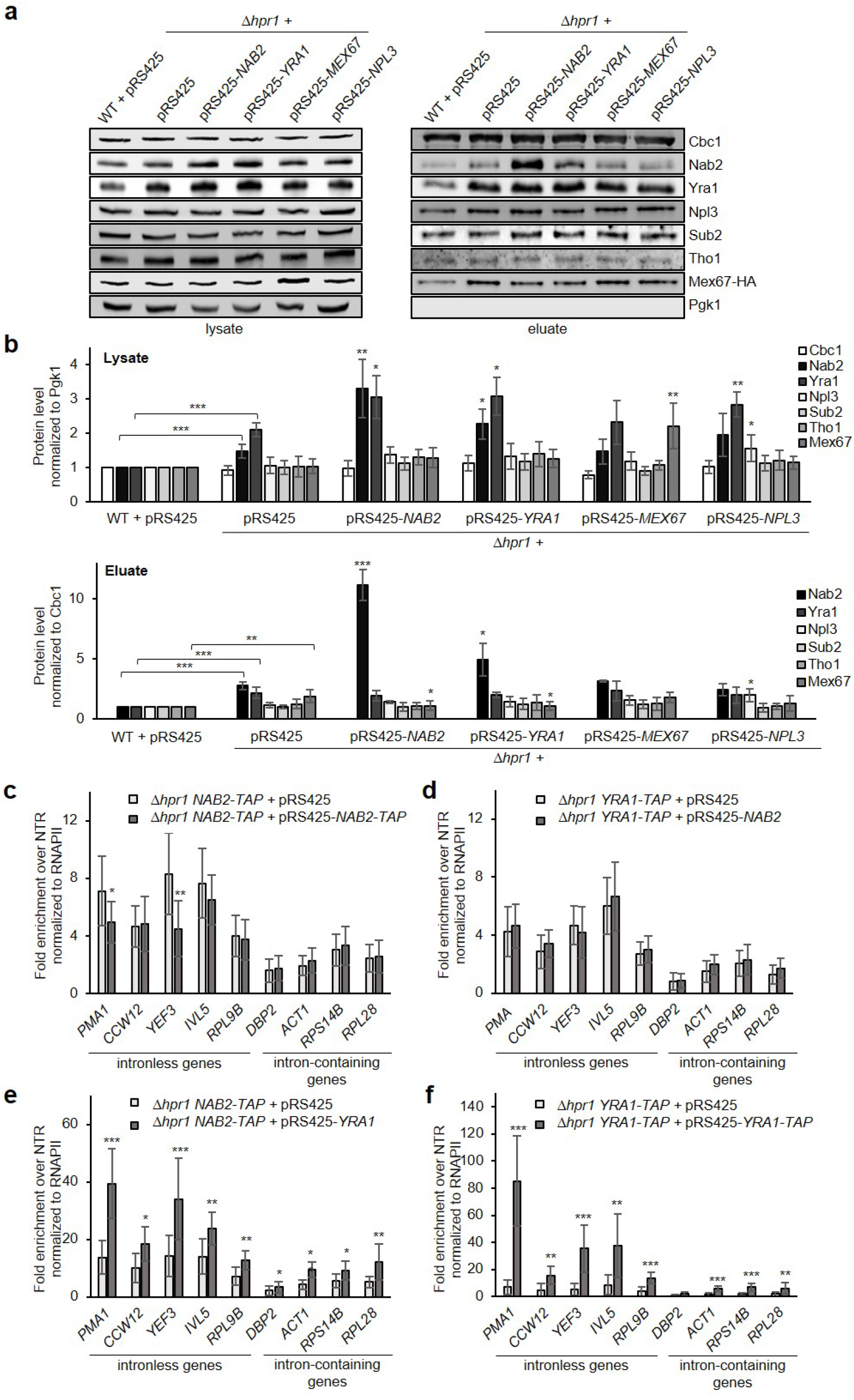
Overexpression of Nab2 or Yra1 in *Δhpr1* cells leads to a further increase of the Nab2 level and a decrease in the Mex67 level in nuclear mRNPs. **a** Western blots of lysates and TEV eluates of nuclear mRNPs obtained by Cbc2-TAP purification, using antibodies against indicated RBPs. **b** Quantification of protein levels in lysates and TEV eluates were normalized to the signal of Pgk1 or Cbc1, respectively. Protein levels for WT cells were set to 1. Asterisks with brackets represent the comparison between WT and *Δhpr1* cells, asterisks without brackets indicate the comparison to *Δhpr1* cells. **c - d** The occupancy of Nab2 (**c**) or Yra1 (**d**) at transcribed genes does not change upon Nab2 overexpression in *Δhpr1* cells. **e - f** The occupancy of Nab2 (**e**) or Yra1 (**f**) increases upon Yra1 overexpression in *Δhpr1* cells. Protein occupancy was assessed by ChIP as in Fig. 1f - h and normalized to the occupancy of RNAPII in the respective strains. Data represents the mean ± SD of at least three independent experiments; *p<0.05; **p<0.01; *** p<0.001.

In order to determine, whether the increased Nab2 level in nuclear mRNPs of *Δhpr1* cells by overexpression of Nab2 or Yra1 is already present at transcribed genes, we assessed the occupancy of Nab2 and Yra1 by ChIP (Fig. 3c-f). When Nab2 is overexpressed in *Δhpr1* cells the occupancy of Nab2 or Yra1 does not change (Fig. 3c, d). Thus, “additional” Nab2 is most likely incorporated into nuclear mRNPs after transcription. In contrast, overexpression of Yra1 in *Δhpr1* cells leads to a higher occupancy of Nab2 as well as of Yra1 (Fig. 3e, f). This increased occupancy correlates with higher levels of Nab2 but not of Yra1 in nuclear mRNPs (Fig. 3a, b). Thus, Nab2 and Yra1 act in an overlapping yet different manner: Nab2 overexpression does not change Nab2 or Yra1 occupancy but results in a higher Nab2 level in nuclear mRNPs. In contrast, overexpression of Yra1 increases the occupancy of Yra1 and Nab2, but yields only elevated levels of Nab2 in nuclear mRNPs, suggesting that Yra1 either recruits Nab2 or stabilizes its interactions for its consecutive loading onto mRNPs. This is consistent with the finding that Yra1 is not absolutely required for nuclear mRNA export and rather acts as a cofactor favoring the interaction between Nab2 and Mex67^20^. In summary, higher levels of Nab2 or Yra1 suppress the mRNA export defect of *Δhpr1* cells and reduce the levels of Mex67 in nuclear mRNPs to WT levels.

### Nuclear pore complex mutants with a nuclear mRNA export defect have higher levels of Nab2, Yra1 and Mex67 in nuclear mRNPs, but are not suppressed by overexpression of Nab2 or Yra1

To examine whether the increased Nab2 and Yra1 level in nuclear mRNPs is a general phenomenon of nuclear mRNA export mutants, we analyzed strains with deletion of *NUP60, NUP133* or *NUP2* coding for nuclear pore proteins or of *SAC3* coding for a THSC/TREX2 component, all of which exhibit a nuclear mRNA export defect (Fig. 4a and ^16^). Interestingly, the Nab2, Yra1 and Mex67 levels in nuclear mRNPs in all four mutants are increased similarly to *Δhpr1* cells (Fig. 4b, c). In contrast, overexpression of Yra1 or Nab2 does not suppress the nuclear mRNA export defect of *Δnup60, Δnup133, Δnup2* and *Δsac3* cells (Supplementary Fig. 3a-d). To determine the potential reason for this lack of suppression we analyzed the composition of nuclear mRNPs for one of these strains, *Δnup60*. As in the *Δhpr1* mutant, overexpression of either Nab2 or Yra1 leads to higher levels of Nab2 in the *Δnup60* mutant (Supplementary Fig. 3e, f). In contrast to *Δhpr1* cells, not only the Nab2, but also the Yra1 level in nuclear mRNPs increases due to overexpression of Nab2 or Yra1 (Supplementary Fig. 4e, f). Importantly, the Mex67 level in nuclear mRNPs is still elevated compared to WT in *Δnup60* cells with Nab2 or Yra1 overexpression (Supplementary Fig. 3e, f), concomitant with the persisting nuclear mRNA export defect. Thus, an mRNA export defect caused by mutation at a later stage of nuclear mRNA export, i.e. at the nuclear pore, apparently cannot be suppressed by overexpression of Nab2 or Yra1.

**Figure 4.**
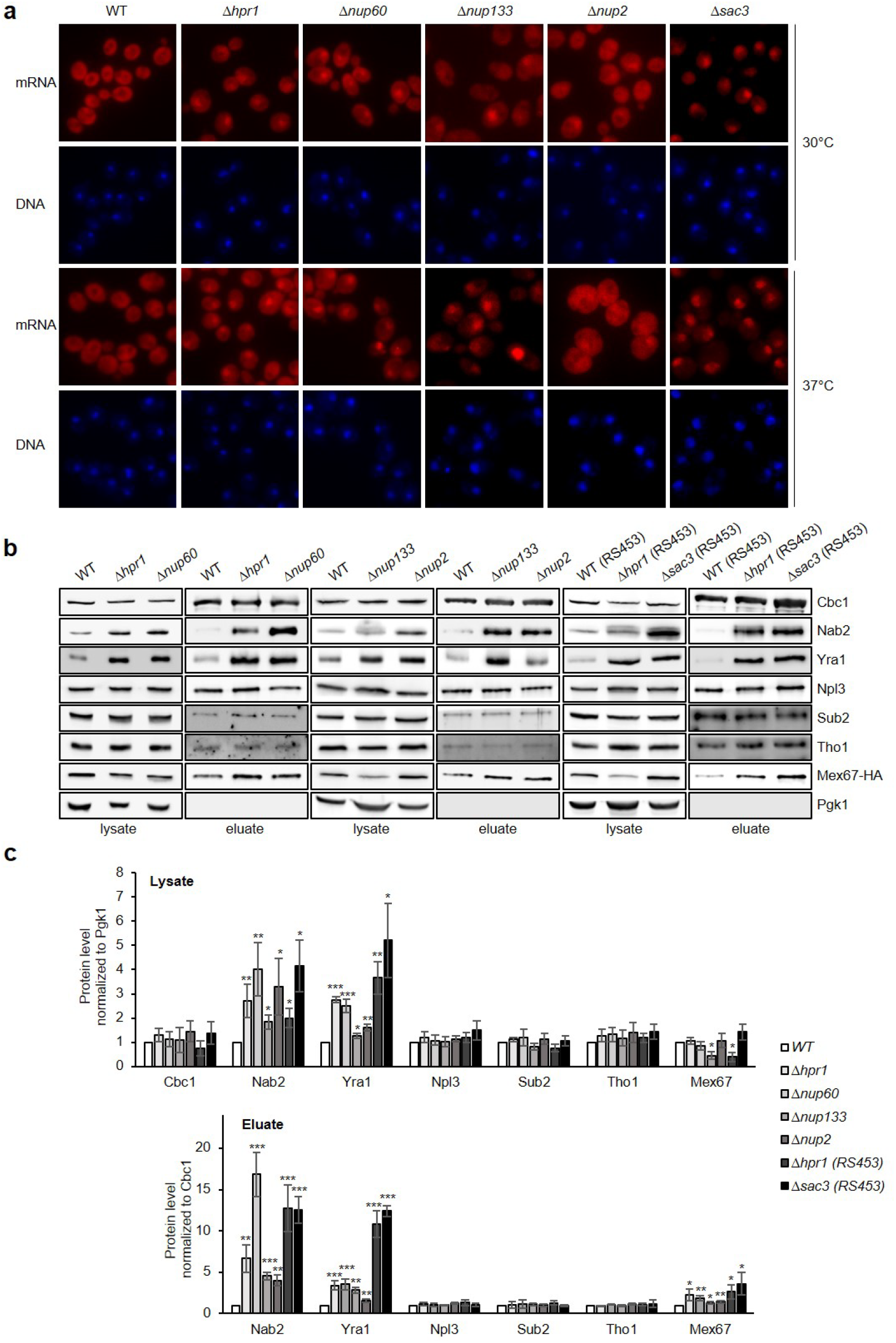
Deletion of *NUP60, NUP133, NUP2* or *SAC3* results in an increase of Nab2, Yra1 and Mex67 levels in nuclear mRNPs. **a** Nuclear mRNA export is impaired in *Δhpr1, Δnup60, Δnup133, Δnup2* and *Δsac3* cells. Poly(A)+ RNA was visualized by FISH at 30°C or after 1 h shift to 37°C with a Cy3-conjugated oligo(dT)50 probe. DNA was stained with DAPI. **b** Western blots of lysates and TEV eluates after Cbc2-TAP purification from WT, *Δnup60, Δnup133, Δnup2* and *Δsac3* cells, using antibodies against the indicated RBPs. **c** Quantification of protein levels in lysates and TEV eluates, normalized to Pgk1 or Cbc1, respectively. Values for WT cells were set to 1. Data represents the mean ± SD of three independent experiments; *p<0.05; **p<0.01; *** p<0.001. Unless otherwise indicated all used strains have a W303 background. As deletion of *SAC3* causes a stronger nuclear mRNA export defect in the RS453 than in the W303 background the *Δsac3* strain, and for comparison the WT and *Δhpr1* strains, has an RS453 background.

### Depletion of Sub2 causes a nuclear mRNA export defect but does not lead to an increased Mex67 level in nuclear mRNPs

Interestingly, the level of the mRNA exporter Mex67 in nuclear mRNPs is increased in *Δhpr1* cells, which have an mRNA export defect, and is reduced to WT level in *Δhpr1* cells by overexpression of Nab2 or Yra1, which represses the nuclear mRNA export defect. Thus, the increase of the Mex67 level in nuclear mRNPs could be either cause or consequence of the nuclear mRNA export defect. As depletion of the nuclear mRNP component Sub2 causes a nuclear mRNA export defect^32^, we determined the composition of nuclear mRNPs after Sub2 depletion. As for the other mRNA export mutants, Nab2 and Yra1 levels in nuclear mRNPs increase (Fig. 5a, b). Furthermore, the Tho1 level increases, whereas the Npl3 level decreases in nuclear mRNPs (Fig. 5a, b). Importantly, the Mex67 level in nuclear mRNPs is unchanged (Fig. 5a, b). Therefore, the elevated Mex67 level in nuclear mRNPs of cells lacking TREX, THSH or nuclear pore proteins is not the result but rather the cause of defective nuclear mRNA export. In addition, higher Nab2 and Yra1 levels in nuclear mRNPs do not necessarily result in an increase in the Mex67 level in nuclear mRNPs. In summary, not only the lack of the mRNA exporter Mex67-Mtr2 and thus its function, but also an increased level of Mex67 in nuclear mRNPs causes a defect in nuclear mRNA export.

**Figure 5.**
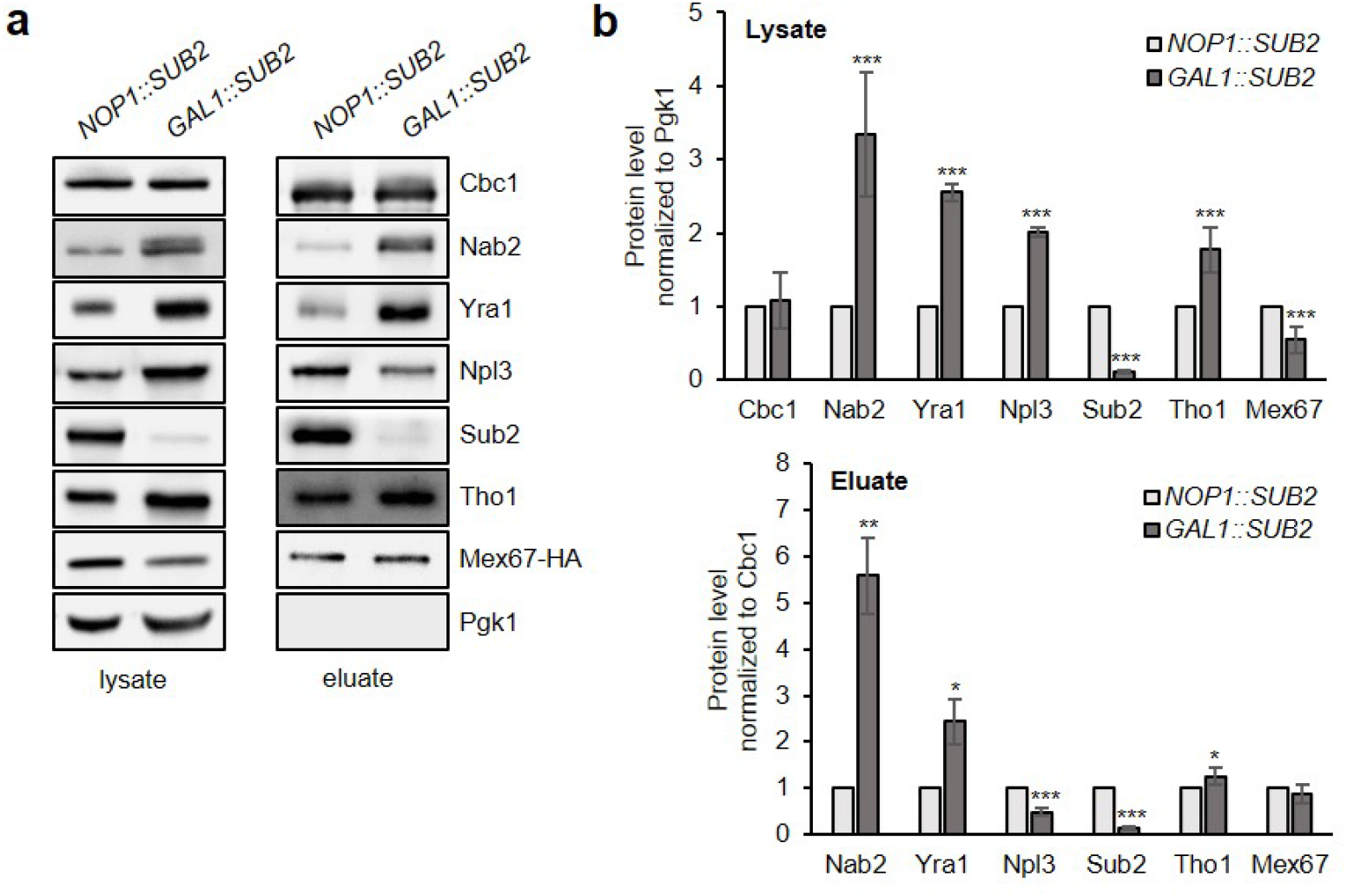
A nuclear mRNA export defect does not always cause an elevated Mex67 level in nuclear mRNPs. **a** Western blots of lysates and TEV eluates after Cbc2-TAP purification from cells expressing Sub2 (*pNOP1::SUB2*) or after depletion of Sub2 (*pGAL1::SUB2*), using antibodies against the indicated RBPs. Cells either expressing Sub2 from a constitutive promoter (*pNOP1::SUB2*) or from a galactose-inducible promoter (*pGAL1::SUB2*) were shifted from galactose-to glucose-containing medium for 20 h to deplete Sub2. **b** Quantification of protein levels of lysates and TEV eluates, normalized to Pgk1 or Cbc1, respectively. Values for WT cells were set to 1. Data represents the mean ± SD of three independent experiments; *p < 0.05; **p < 0.01; *** p < 0.001.

## Discussion

Assembly of the mRNA into an mRNP in the nucleus is an essential prerequisite for nuclear mRNA export. Nevertheless, how the many different mRNP components bind to the mRNA as well as the stoichiometry and the architecture of a nuclear mRNP are still unknown. Furthermore, many if not all mRNP components have several functions in gene expression. Among other things, it is unclear why different nuclear mRNA export adaptors exist and what their precise functions are. In *S. cerevisiae*, there are four adaptors: Hpr1, Nab2, Yra1 and Npl3. Interestingly, cells compensate the lack of Hpr1, which leads to disassembly of the THO complex, by increasing total levels as well as the levels of Nab2 and Yra1 in nuclear mRNPs. This might be an attempt to increase nuclear mRNA export in *Δhpr1* cells rendering these cells viable despite a defect in mRNP assembly and consequently in nuclear mRNA export. The absence of Hpr1 and consequently of the large THO complex on the mRNA could expose larger mRNA stretches to Nab2 and Yra1. Nab2 and Yra1 might – partially – compensate for the loss of THO by binding to the mRNA and inducing its compaction^33,34^.

Interestingly, overexpression of Nab2 or Yra1 in *Δhpr1* cells with already increased Nab2 and Yra1 levels suppresses their growth as well as nuclear mRNA export defect. This suggests that Nab2 and Yra1 might have similar functions. Consistently, Yra1 is dispensable for nuclear mRNA export in *Drosophila* and *Caenorhabditis elegans*^20,35,36^. Moreover, Yra1 might recruit Nab2 to be loaded onto the mRNP, because overexpression of Yra1 in *Δhpr1* cells leads to increased occupancy of both proteins, Yra1 and Nab2. Interestingly, the mRNA exporter Mex67-Mtr2 interacts directly with both adaptor proteins, and Yra1 enhances the interaction between Nab2 and Mex67, which is supported by the fact that Yra1 is not essential in cells overexpressing Nab2 or Mex67^18,20^. Taken together, in wild-type cells Nab2 and Yra1 might form a complex that recruits the mRNA export adaptor Mex67-Mtr2 to the nuclear mRNP.

In contrast, the adaptor protein Npl3 seems to be different from Nab2 and Yra1. Npl3 levels in nuclear mRNPs do not increase in *Δhpr1* cells, and Npl3 overexpression does not suppress the growth and nuclear mRNA export defects of *Δhpr1* cells. Consistently, Nab2 and Npl3 preferentially associate with a distinct set of mRNAs as Npl3 largely associates with transcripts coding for ribosomal proteins and other highly expressed transcripts, while Nab2 associates with transcripts coding for proteins required for transcription^37^. Thus, either the mechanistic function or the subset of bound mRNAs might be different between Nab2 / Yra1 and Npl3.

Importantly, we were not able to identify other defects in *Δhpr1* cells such as transcription, transcriptional readthrough, poly(A) tail length, splicing or pre-mRNA leakage that are suppressed by overexpression of Nab2 or Yra1. As a proxy for transcription, RNAPII occupancy is reduced in *Δhpr1* cells (Supplementary Fig. 2). However, this transcriptional defect in *Δhpr1* cells is not suppressed by overexpression of Nab2 or Yra1 as RNAPII occupancy does not increase (data not shown). Furthermore, *Δhpr1* cells accumulate R loop structures, composed of DNA-RNA hybrids and the displaced single-stranded DNA strand, which contribute to genomic instability^38^. However, this higher amount of R loops in *Δhpr1* cells is not suppressed by overexpression of Nab2 or Yra1 (data not shown). Likewise, although loss of Nab2 or Npl3 results in global transcriptional readthrough^13,39^, overexpression of Nab2 or Yra1 does not suppress readthrough at the tested examplary genes in *Δhpr1* cells (data not shown). Nab2 is a nuclear poly(A) binding protein regulating poly(A) tail length, and a nuclear mRNA export defect often causes hyperadenylation^40,41^. However, the poly(A) tails of bulk RNA as well as of single examplary transcripts are of wild-type length in *Δhpr1* cells, and their length does not change due to overexpression of Nab2 or Yra1 (data not shown). Furthermore, splicing of a reporter transcript rather increases in *Δhpr1* cells, most likely reflecting their longer dwell time in the nucleus due to the mRNA export defect (data not shown). Last but not least, leakage of an intron-containing reporter transcript to the cytoplasm is increased in *Δhpr1* cells. This defect is also not suppressed by Nab2 or Yra1 overexpression (data not shown). Thus, overexpression of Nab2 or Yra1 in *Δhpr1* cells exclusively alleviates the nuclear mRNA export defect and, most likely as a consequence, their growth defect.

The adaptor proteins Nab2 and Yra1 recruit the mRNA exporter Mex67-Mtr2 to the nuclear mRNP, and Mex67-Mtr2 export the mRNP though the nuclear pore complexes (Fig. 6a). Loss of the THO complex leads to higher levels of the mRNP components Nab2, Yra1 and Mex67 in nuclear mRNPs compared to wild-type cells and a nuclear mRNA export defect (Fig. 1 and Fig. 6b). When nuclear mRNA export is impaired by deletion of nuclear pore or THSC components, the mRNP composition changes in a similar way as in *Δhpr1* cells (Fig. 6b). Importantly, the higher Mex67 level in nuclear mRNPs is most likely not a consequence of the nuclear mRNA export defect as depletion of the mRNP component Sub2 likewise causes a nuclear mRNA export defect but does not change the Mex67 level in nuclear mRNPs (Fig. 5). Importantly, overexpression of Nab2 or Yra1 not only suppresses the nuclear mRNA export defect of *Δhpr1* cells but also decreases the elevated Mex67 level in nuclear mRNPs to wild-type levels (Fig. 2b, Fig. 3a, b and Fig. 6c). Consistently, overexpression of Mex67 in *Δhpr1* cells neither suppresses the mRNA export defect nor changes the composition of nuclear mRNPs (Fig. 2b and Fig. 3a, b). The fact that suppression of the nuclear mRNA export defect coincides with a reduced Mex67 level in nuclear mRNPs (compared to the composition of nuclear mRNPs in *Δhpr1* cells, which display a nuclear mRNA export defect) is surprising as Mex67-Mtr2 – according to the current model – needs to be recruited to nuclear mRNPs for their export. Here, we broaden this model by showing that an increased level of Mex67-Mtr2 also causes a nuclear mRNA export defect.

**Figure 6.**
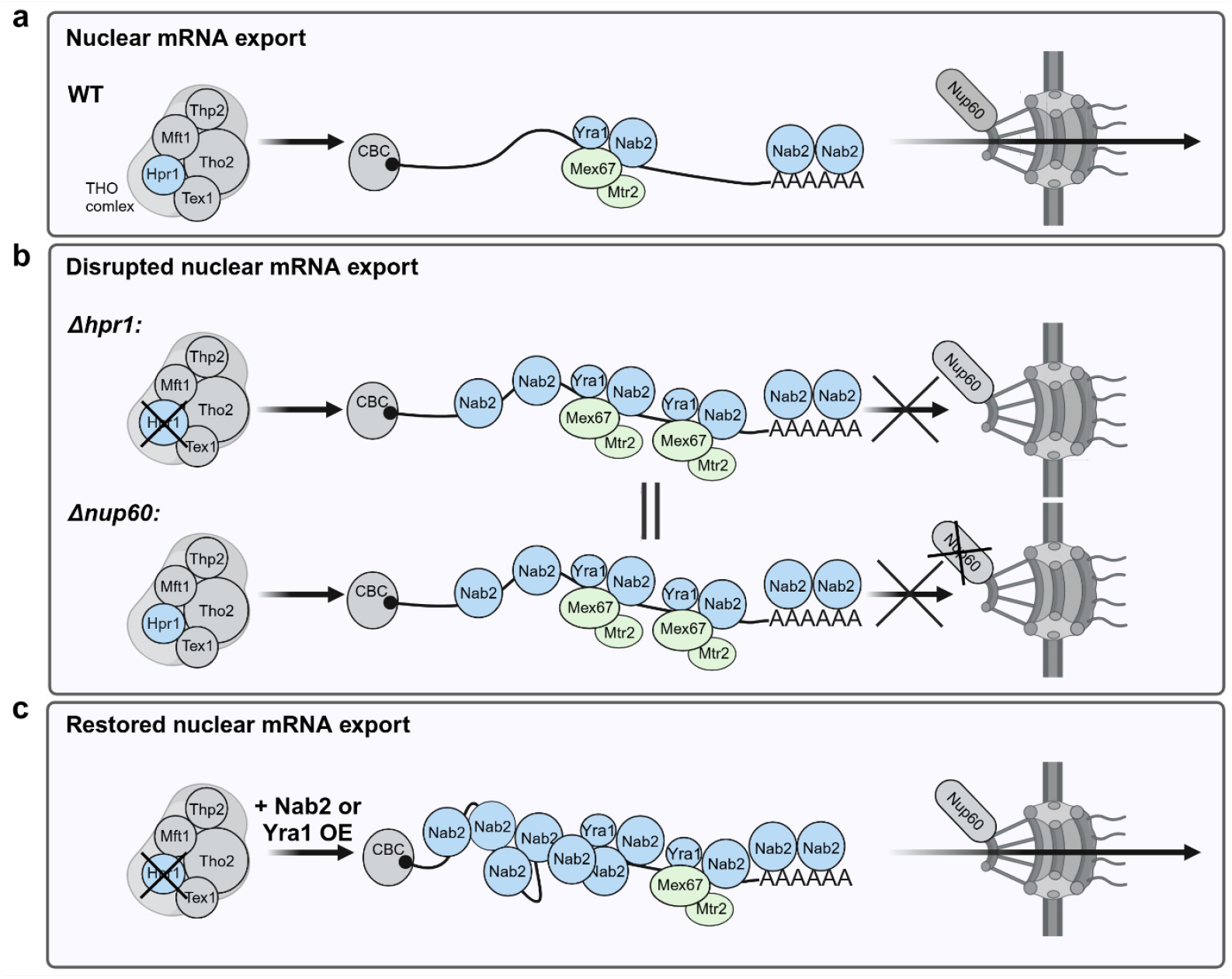
Nuclear mRNA export depends on a balanced level of the mRNA exporter Mex67-Mtr2 in nuclear mRNPs. **a** Nuclear mRNP assembly is one prerequisite for nuclear mRNA export. In a WT strain, the adaptor proteins Hpr1, Nab2, Yra1 and Npl3 recruit the mRNA exporter Mex67-Mtr2 to the nuclear mRNP for nuclear mRNA export. Npl3 and all other nuclear mRNP components are not shown for clarity. **b** Loss of Hpr1 and thus the THO complex causes increased levels of Nab2, Yra1 and Mex67 in nuclear mRNPs and a nuclear mRNA export defect. Disruption of mRNA export at the nuclear pore by deletion of *NUP60* or other nuclear pore components such as *NUP133, NUP2* or *SAC3* results in similar changes in the composition of nuclear mRNPs and a nuclear mRNA export defect. **c** Overexpression of Nab2 or Yra1 in *Δhpr1* cells further increases the Nab2 level and decreases the Mex67 level to WT in nuclear mRNPs and restores nuclear mRNA export. Therefore, a wild-type level of Mex67 is required for efficient nuclear mRNA export.

Indeed, the level of Mex67-Mtr2 in nuclear mRNPs is likely tightly regulated as overexpression of Mex67-Mtr2 in wild-type cells neither leads to a higher level of Mex67 or any other mRNP component in nuclear mRNPs nor to a nuclear mRNA export defect (Supplementary Fig. 4 a-c). Similarly, overexpression of Nab2, Yra1 or Nab2 and Yra1 in wild-type cells does not cause a nuclear mRNA export defect (Supplementary Fig. 4c). Interestingly, even though overexpression of Nab2 in wild-type cells does not cause a nuclear mRNA export defect, Nab2 as well as Yra1 levels in nuclear mRNPs increase (Supplementary Fig. 4 d, e). However, the level of Mex67 in nuclear mRNPs does not change – in contrast to *Δhpr1* cells, showing that higher levels of Nab2 and Yra1 alone do not lead to an increased Mex67 level and thus not a nuclear mRNA export defect (Supplementary Fig. 4 d, e). In summary, wild-type levels of Mex67-Mtr2 in nuclear mRNPs are needed for efficient nuclear mRNA export.

## Methods

### Strains, plasmids and primers

Yeast strains, plasmids and primers are listed in Supplementary Tables 1-3, respectively.

### Purification of nuclear mRNPs

Nuclear mRNPs were isolated by purification of endogenously TAP-tagged Cbc2, a subunit of CBC, as described in ^12^ with some modifications. Briefly, 2 l of yeast culture were harvested at OD_600_ 3.5 and lysed by cryogenic grinding (6870D Freezer/Mill, SPEX Sample Prep). The grindates were mixed with 10 ml lysis buffer (50 mM HEPES-KOH pH 7.5, 100 mM KCl, 1.5 mM MgCl_2_, 0.15% NP-40, 1 mM DTT, 1x protease inhibitor), pre-cleared by centrifugation at 3,500 g for 12 min and spun at 165,000 g for 1 h at 4°C. The supernatants were incubated with 500 μl of IgG Sepharose 6 Fast Flow beads (GE Healthcare) for 1.5 h at 4°C. The beads were washed with 15 ml wash buffer (50 mM HEPES-KOH pH 7.5, 200 mM KCl, 1.5 mM MgCl_2_, 0.15% NP-40, 0.5 mM DTT) and incubated with TEV protease for 1 h at 16°C to elute bound proteins. Proteins in lysate and eluate were analyzed by Western blotting using the respective primary antibodies against the protein of interest, followed by incubation with a horse radish peroxidase-coupled secondary antibody. Chemiluminescent signals were detected with CheLuminate-HRP ECl solution (Applichem) using a ChemoCam Imager (Intas). Protein intensities were quantified using ImageJ software.

### RNA immunoprecipitation

RNA immunoprecipitation was performed as described in ^12^. 400 ml of cells expressing the endogenously TAP-tagged protein of interest were harvested at OD_600_0.8, resuspended in 1 ml RNA-IP buffer (25 mM Tris-HCl pH 7.5, 150 mM NaCl, 2 mM MgCl_2_, 0.2% Triton X-100, 0.5 mM DTT, 2x protease inhibitor) and lysed with glass beads using FastPrep-24 5G device (3x for 20 sec at 6 m/s). Cell debris was cleared by centrifugation for 15 min at 16,000 g at 4°C. 900 μl of the lysate were incubated with 660 units DNaseI for 30 min on ice (Input). 35 μl IgG-coupled Dynabeads (Thermo Scientific) were added to the lysate and incubated for 3 h at 4°C on a rotating wheel. The beads were washed 8x with 1 ml RNA-IP buffer. 1 ml of TRIzol was added to the beads for RNA extraction (IP). Isolated RNA from Input and IP samples was reverse-transcribed for subsequent qPCR analysis using the Applied Biosystems StepOnePlus cycler. Standard curves were used to determine qPCR efficiency (E). mRNA enrichment was calculated relative to the untagged negative control (nc) according to the formula: *[E^(Ct IP – Ct Input)nc]/[E^(Ct IP-Ct Input)gene]*.

### Chromatin immunoprecipitation

Chromatin immunoprecipitation (ChIP) was performed as described in ^42^. 100 ml of cell culture were grown to OD_600_ 0.8 and cross-linked with 1% formaldehyde for 20 min. Cells were washed, resuspended in 800 μl low-salt buffer (50 mM HEPES-KOH pH 7.5, 150 mM NaCl, 1 mM EDTA, 1% Triton X-100, 0.1% SDS, 0.1% Sodium Deoxycholat + 1x protease inhibitor) and lysed with glass beads in a FastPrep-24 5G device (2x 45 sec at 6 m/s). Chromatin was fragmented to 200 - 250 bp by sonication with a Bioruptor UCD-200 (Diagenode) 3x for 15 min (30 s ON / 30 s OFF; “HIGH” mode). Lysates were cleared by centrifugation for 15 min at 16,000 g at 4°C (Input). To immunoprecipitate a TAP-tagged protein, the lysate was incubated with 15 μl IgG-coupled Dynabeads M-280 (Thermo Scientific) for 2.5 h at RT. For RNAPII ChIP, the lysate was incubated with 4 μl 8WG16 antibody (BioLegend) for 1.5 h followed by addition of 15 μl Dynabeads Protein G (Thermo Scientific) for 1 h. Beads were washed with 800 μl of buffer (2x low-salt buffer, 3x high-salt buffer (500 mM NaCl), 3x TLEND and 2x with 1x TE pH 8.0) and DNA-protein complexes were eluted (IP). Input and IP samples were incubated with proteinase K for 2 h at 37°C followed by reversal of the crosslinks for 14 h at 65°C. DNA was purified with the PCR NucleoSpin® Gel and PCR Clean-up-kit (Macherey-Nagel) and used for qPCR analysis. Standard curves were used to determine the qPCR efficiency (E). As a negative control, a non-transcribed region (NTR) on chromosome V (174131–174200) was used. Protein occupancy was calculated as the enrichment of the sequence for the gene of interest relative to the NTR using the formula: *[E^(Ct IP – Ct Input)NTR]/[E^(Ct IP – Ct Input)Gene]*.

### Dot spot assay

Yeast cells were resuspended in 2 ml water. The cell suspensions were diluted to the same OD_600_ of 0.09 for each strain (1^st^ spot). 5 μl of 10-fold serial dilutions were spotted on selective media plates. Cells were grown at 25°C, 30°C and 37°C for 2 days, and at 16°C for up to 6 days before imaging.

### Fluorescence *in situ* hybridization

The cellular localization of poly(A)+ RNA was determined by fluorescence *in situ* hybridization (FISH) performed as described previously ^12^ with some modifications. Cells were grown to an OD_600_ 0.8 at 30°C or shifted to 37°C for 1 h before crosslinking with 4% formaldehyde for 90 min. Washed cells were incubated with 100 μg 100T zymolase for 30 min at 30°C to digest the cell walls. Obtained spheroplasts were applied to polylysine-coated coverslips. The adherent cells were incubated with 200 μl 2x SSC buffer for 10 min, followed by incubation with 12 μl of prehybridization buffer (50% formamide, 10% dextran sulphate, 125 μg/ml of *Escherichia coli* tRNA, 500 μg/ml herring sperm DNA, 4x SSC, 0.02% polyvinyl pyrrolidone, 0.02% BSA, 0.02% Ficoll-40) for 1 h at 37°C in a humid chamber. 0.75 μl of 1 pmol/μl oligo(dT)50-Cy3 probe was added and incubated at 37°C overnight in a humid chamber. After hybridization, coverslips with cells were washed with 3 ml 0.5x SSC for 30 min on a rocking shaker, air-dried and mounted with ROTI® Mount FluorCare DAPI (Carl Roth) on microscope slides. Cells were imaged using a Deltavision Ultra High-Resolution Microscope (Cytiva).

## Statistical analysis

All experiments were performed with at least three independent biological replicates. Data is presented as mean ± standard deviation (error bars). Asterisks indicate statistical significance (Student’s *t*-test; \**P* ≤ 0.05; \*\**P* ≤ 0.01; \*\*\**P* ≤ 0.001).

## Acknowledgements

We thank Vera Bettenworth for critical reading of the manuscript and all other members of the KS and the Cornelia Kilchert group for fruitful discussions. We are also grateful to Salome Barbadkadze for preliminary ChIP experiments. The antibody directed against Cbp80 was a kind gift of Dirk Görlich (MPI Goettingen, Germany)^43^.

