## Supplemental Data for "An elevated level of the mRNA exporter Mex67-Mtr2 in nuclear mRNPs impairs nuclear mRNA export"

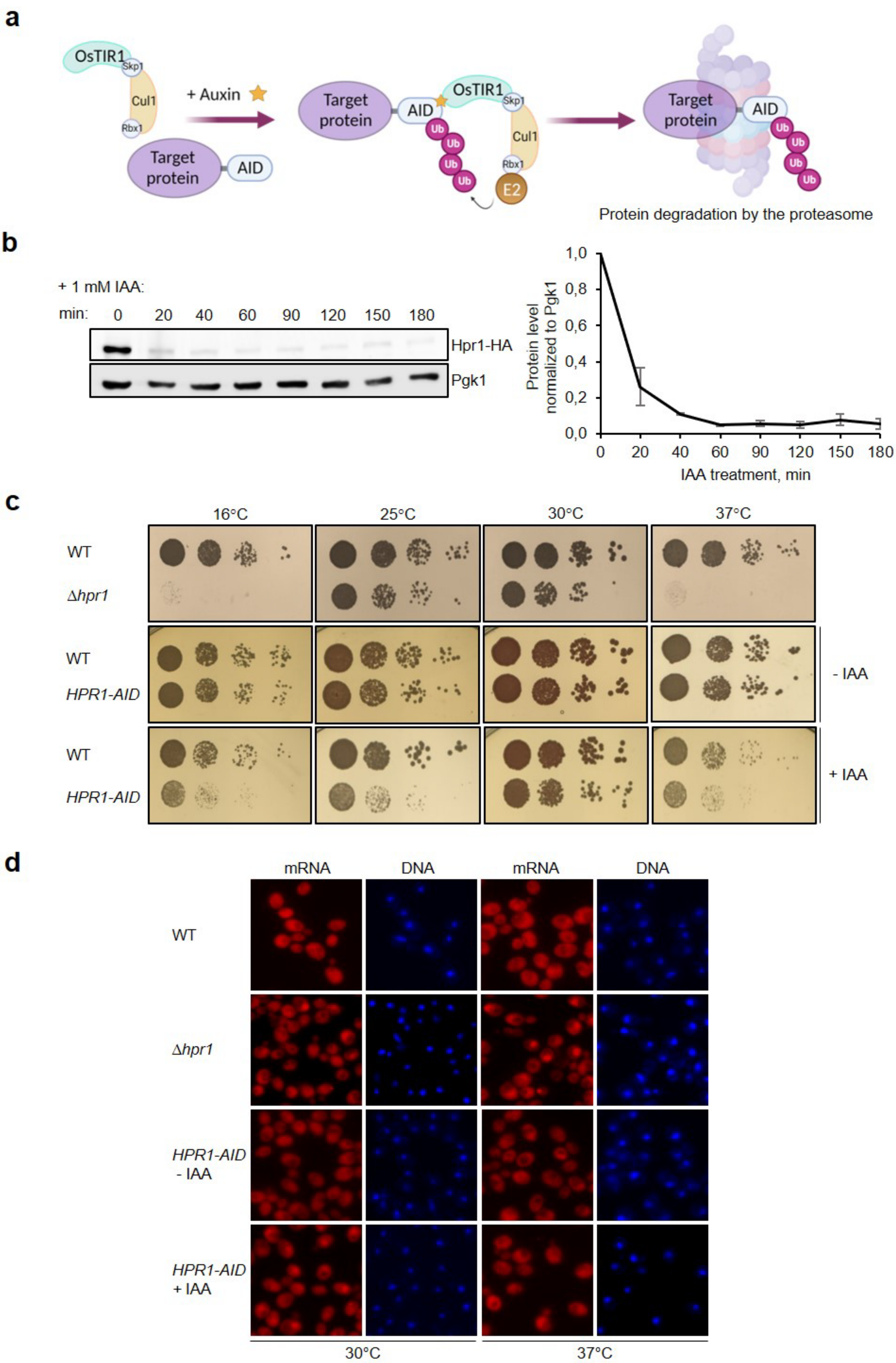
a
OsTIR1
Skp1
Cul1
Rbx1
Target protein
AID
+ Auxin
Target protein
AID
OsTIR1
Skp1
Cul1
Rbx1
E2
Ub
Target protein
AID
Ub
Protein degradation by the proteasome
b
+ 1 mM IAA:
min: 0 20 40 60 90 120 150 180
Hpr1-HA
Pgk1
Protein level normalized to Pgk1
IAA treatment, min
c
16°C
25°C
30°C
37°C
WT
Δhpr1
WT
HPR1-AID
- IAA
WT
HPR1-AID
+ IAA
d
mRNA
DNA
mRNA
DNA
WT
Δhpr1
HPR1-AID - IAA
HPR1-AID + IAA
30°C
37°C

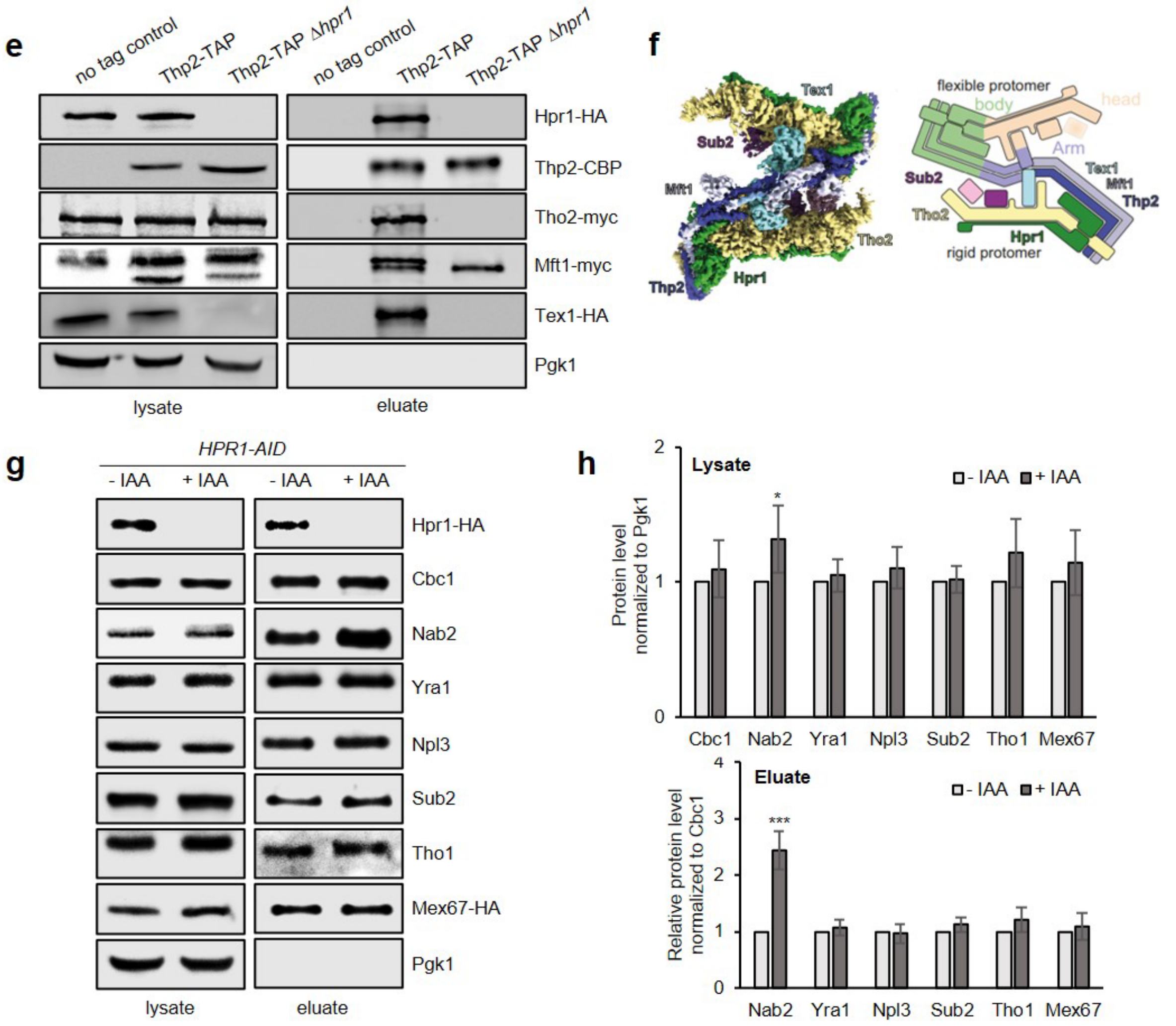


**Supplementary Fig. 1. Depletion of Hpr1 impairs cell growth and nuclear mRNA export and leads to increased Nab2 level in nuclear mRNPs. a** Scheme of the auxin-inducible degron (AID) system (modified from [1]). Addition of auxin promotes the interaction between SCF E3 ligase complex subunit OsTIR1 and the AID tag of the target protein. This causes the recruitment of an E2 ligase that ubiquitinates the AID tag and thus targets the tagged protein for degradation by the proteasome. **b** Hpr1-AID is depleted after addition of auxin. Western blots of whole cell extracts after incubation of *HPR1-AID-HA* cells with 1 mM auxin (IAA) at different time points. Hpr1-AID-HA and Pgk1, which served as loading control, were detected with the respective antibodies. Hpr1 levels were quantified and normalized to Pgk1 levels. Data represents the mean ± standard deviation (SD). **c** Deletion or depletion of Hpr1 causes a growth defect. Ten-fold serial dilutions of WT and *Δhpr1* cells and of cells expressing Hpr1-AID were spotted onto YPD plates without (- IAA) or with 1 mM auxin (+ IAA), respectively, and grown at the indicated temperatures. **d** Deletion or depletion of Hpr1 causes a nuclear mRNA export defect. The localization of poly(A)+ RNA was determined at 30°C or after 1 h shift to 37°C of WT and *Δhpr1* cells and of *HPR1-AID* cells incubated without (- IAA) or with 1 mM auxin (+ IAA) for 1 h. Poly(A)+ RNA was visualized with oligo(dT)50-Cy3, DNA was stained with DAPI. **e** Deletion of *HPR1* leads to disintegration of the THO complex. Pull-down of the THO complex via Thp2-TAP in WT and Δ*hpr1* cells. Western blots of lysates and TEV eluates after cleavage of the TAP tag, using antibodies against tagged components of the THO complex. Note, that the total amount of Tex1 decreases by deletion of *HPR1*. **f** Cryo-EM structure of the reconstituted THO-Sub2 complex (left) and its scheme (right) reproduced from [2]. **g** Acute depletion of Hpr1 causes an elevated Nab2 level in nuclear mRNPs. Western blots of lysates and TEV eluates after Cbc2-TAP purification from cells expressing Hpr1-AID without auxin treatment (-IAA) or with 1 mM auxin treatment (+ IAA) for 1 h using antibodies against the indicated RBPs. **h** Quantification of the protein levels in lysates (upper panel) and TEV eluates (lower panel) normalized to Pgk1 or Cbc1 levels, respectively. Values for WT cells were set to 1. Data represents the mean ± SD from at least three independent experiments; *$p<0.05$; *** $p<0.001$.

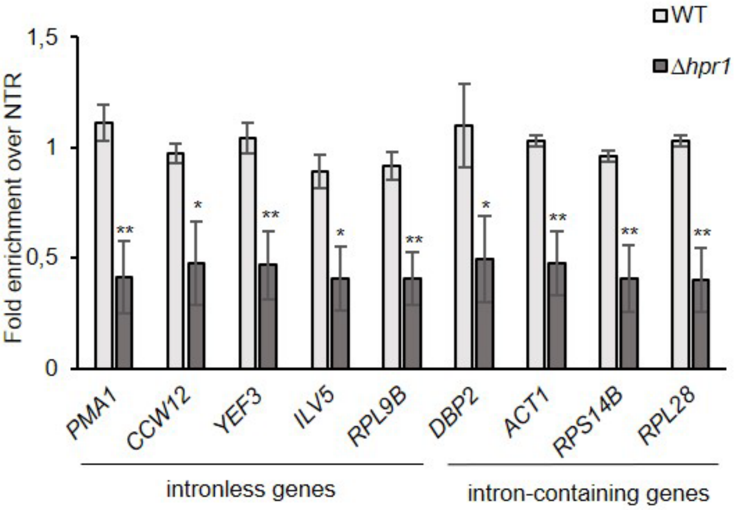


**Supplementary Fig. 2. RNAPII occupancy decreases in Δ*hpr1* cells.**
The occupancy of the RNAPII subunit Rpb1 at transcribed genes was assessed by chromatin immunoprecipitation (ChIP) using the antibody 8WG16, which recognizes the non-phosphorylated CTD. The occupancy was calculated as the enrichment of protein at the exemplary genes shown relative to its presence at a non-transcribed region (NTR, 174131–174200 on chr. V) and was set to 1 for the WT strain. Data represents the mean ± SD from at least three independent experiments; *$p<0.05$; **$p<0.01$.

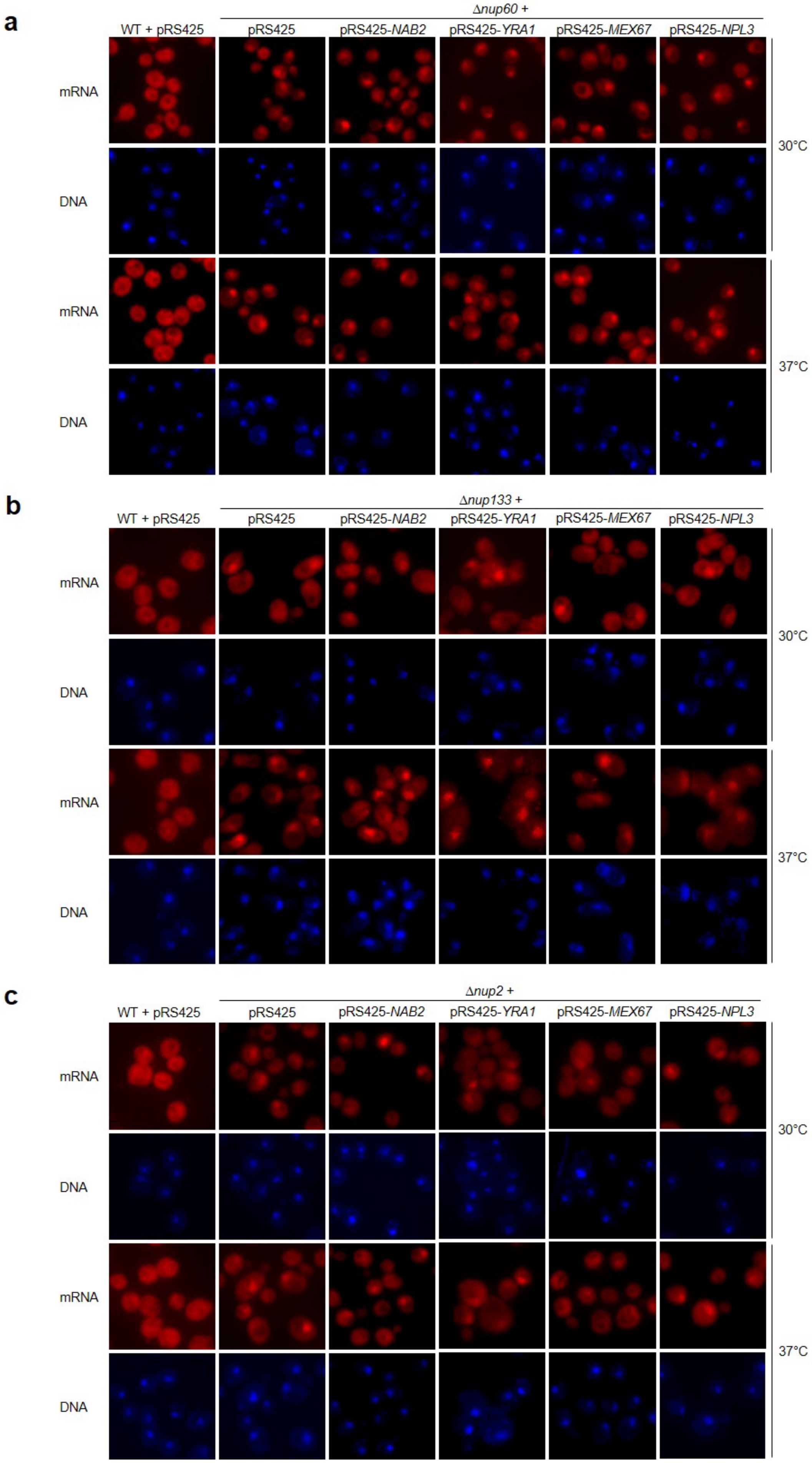
a
Δnup60 +
WT + pRS425
pRS425
pRS425-NAB2
pRS425-YRA1
pRS425-MEX67
pRS425-NPL3
mRNA
DNA
mRNA
DNA
30°C
37°C
b
Δnup133 +
WT + pRS425
pRS425
pRS425-NAB2
pRS425-YRA1
pRS425-MEX67
pRS425-NPL3
mRNA
DNA
mRNA
DNA
30°C
37°C
c
Δnup2 +
WT + pRS425
pRS425
pRS425-NAB2
pRS425-YRA1
pRS425-MEX67
pRS425-NPL3
mRNA
DNA
mRNA
DNA
30°C
37°C

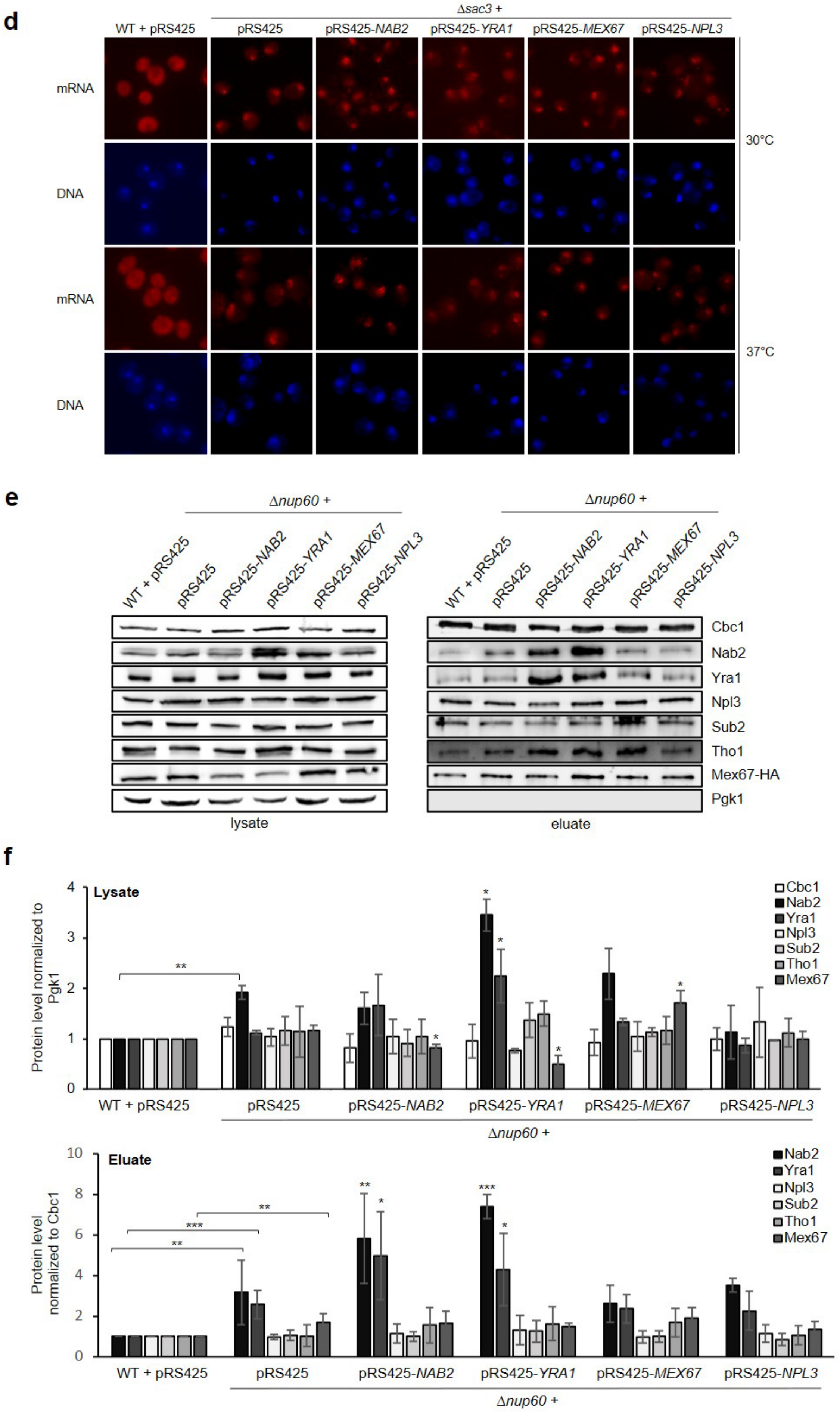

d
Δsac3 +
WT + pRS425
pRS425
pRS425-NAB2
pRS425-YRA1
pRS425-MEX67
pRS425-NPL3
mRNA
DNA
mRNA
DNA
30°C
37°C
e
Δnup60 +
Δnup60 +
WT + pRS425
pRS425
pRS425-NAB2
pRS425-YRA1
pRS425-MEX67
pRS425-NPL3
Cbc1
Nab2
Yra1
Npl3
Sub2
Tho1
Mex67-HA
Pgk1
lysate
eluate
f
Lysate
Protein level normalized to Pgk1
Cbc1
Nab2
Yra1
Npl3
Sub2
Tho1
Mex67
WT + pRS425
pRS425
pRS425-NAB2
pRS425-YRA1
pRS425-MEX67
pRS425-NPL3
Δnup60 +
Eluate
Protein level normalized to Cbc1
Nab2
Yra1
Npl3
Sub2
Tho1
Mex67
WT + pRS425
pRS425
pRS425-NAB2
pRS425-YRA1
pRS425-MEX67
pRS425-NPL3
Δnup60 +

**Supplementary Fig. 3. Overexpression of Nab2, Yra1, Mex67 or Npl3 does not suppress the nuclear mRNA export defect of *Δnup60*, *Δnup133, Δnup2* or *Δsac3* cells and does not decrease Mex67 levels in nuclear mRNPs to WT.**

**a-d** Deletion of *NUP60*, *NUP133*, *NUP2* or *SAC3* causes a nuclear mRNA export defect. Poly(A)+ RNA was visualized by fluorescence in situ hybridization (FISH) with an oligo(dT)50-Cy3 probe at 30°C or after 1 h shift to 37°C in the indicated strains. DNA was stained with DAPI. **e and f** Overexpression of Nab2 or Yra1 in *Δnup60* cells leads to higher levels of Nab2 and Yra1 in nuclear mRNPs but does not decrease the level of Mex67 to WT levels. **e** Representative Western blots of lysates and TEV eluates after Cbc2-TAP purification in *Δnup60* cells and *Δnup60* cells overexpressing Nab2, Yra1, Mex67 or Npl3, using antibodies against the indicated RBPs. **f** Quantification of the protein levels in lysates and TEV eluates, normalized to the signal of Pgk1 or Cbc1, respectively. Protein levels of WT cells were set to 1. Asterisks with brackets represent the comparison between WT and *Δnup60* cells, asterisks without brackets indicate the comparison to *Δnup60* cells.

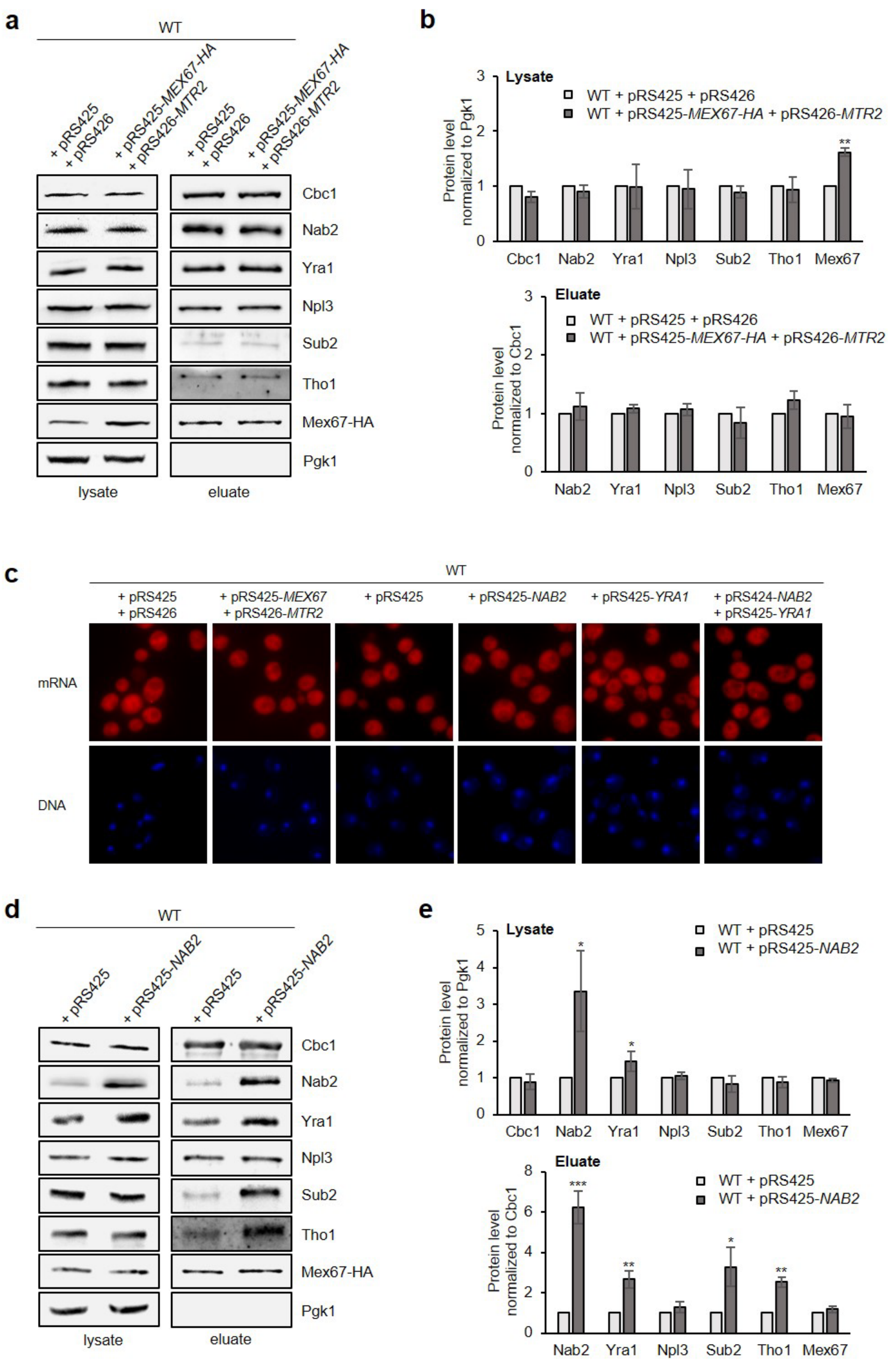
a
WT
+ pRS425 + pRS426
+ pRS425-MEX67-HA + pRS426-MTR2
Cbc1
Nab2
Yra1
Npl3
Sub2
Tho1
Mex67-HA
Pgk1
lysate
eluate
b
Lysate
WT + pRS425 + pRS426
WT + pRS425-MEX67-HA + pRS426-MTR2
Protein level normalized to Pgk1
Cbc1 Nab2 Yra1 Npl3 Sub2 Tho1 Mex67
Eluate
Protein level normalized to Cbc1
Nab2 Yra1 Npl3 Sub2 Tho1 Mex67
c
WT
+ pRS425 + pRS426
+ pRS425-MEX67 + pRS426-MTR2
+ pRS425
+ pRS425-NAB2
+ pRS425-YRA1
+ pRS424-NAB2 + pRS425-YRA1
mRNA
DNA
d
WT
+ pRS425
+ pRS425-NAB2
Cbc1
Nab2
Yra1
Npl3
Sub2
Tho1
Mex67-HA
Pgk1
lysate
eluate
e
Lysate
WT + pRS425
WT + pRS425-NAB2
Protein level normalized to Pgk1
Cbc1 Nab2 Yra1 Npl3 Sub2 Tho1 Mex67
Eluate
Protein level normalized to Cbc1
Nab2 Yra1 Npl3 Sub2 Tho1 Mex67

**Supplementary Fig. 4. Higher Nab2 and Yra1 levels do not result in an increased Mex67 level in nuclear mRNPs nor in a nuclear mRNA export defect. a** Western blots of lysates and TEV eluates of a Cbc2-TAP purification from WT cells or cells overexpressing Mex67-HA and Mtr2, using antibodies against the indicated RBPs. **c** Quantification of protein levels in lysates (upper panel) and TEV eluates (lower panel) normalized to levels of Pgk1 or Cbc1, respectively. Values for WT cells were set to 1. Data represents the mean ± SD of at least three independent experiments; **p <0.01. **c** Overexpression of Mex67 and Mtr2, Nab2, Yra1 or Nab2 and Yra1 does not cause a nuclear mRNA export defect. FISH of WT cells or cells overexpressing Mex67 and Mtr2, Nab2, Yra1 or Nab2 and Yra1 grown at 30°C. poly(A)+ RNA was detected using oligo(dT)50-Cy3 probes, DNA was stained with DAPI. **d** Western blots of lysates and TEV eluates of a Cbc2-TAP purification from WT cells or cells overexpressing Nab2, using antibodies against the indicated RBPs. **e** Quantification of the protein levels in lysates (upper panel) and TEV eluates (lower panel) as in b. Data represents the mean ± SD of at least three independent experiments; *p<0.05; **p <0.01; *** p<0.001.

**Supplementary Table 1. Yeast strains**

| Strain | Genotype | Reference |
|---|---|---|
| W303 | MATa, ura3-1, trp1-1, his3-11,15, leu2-3,112, ade2-1, can1-100, GAL+ | 3 |
| *Δhpr1* | Mata; ura3-1; ade2-1; his3-11,5; trp1-1; leu2-3,112; can1-100; hpr1::HIS3 | R. Rothstein |
| *CBC2-TAP MEX67-6xHA* | MATa; ura3-1; trp1-1; his3-11,15; leu2-3,112; ade2-1; can1-100; GAL+; CBC2-CBP-TEV-protA::TRP1; MEX67-HA::kanMX | This study |
| *CBC2-TAP MEX67-6xHA Δhpr1* | MATa; ura3-1; trp1-1; his3-11,15; leu2-3,112; ade2-1; can1-100; GAL+; hpr1::HIS3; CBC2-CBP-TEV-protA::TRP1; MEX67-HA::kanMX | This study |
| *osTIR HPR1-AID CBC2-TAP MEX67-6xHA* | MATa; ura3-1; trp1-1; his3-11,15; leu2-3,112; ade2-1; can1-100; GAL+; URA3:::osTIR Hpr1-AID-6HA::Hyg; CBC2-CBP-TEV-protA::TRP1; Mex67-6HA::KanMX4 | This study |
| *NAB2-TAP* | MATa; ura3-1; trp1-1; his3-11,15; leu2-3,112; ade2-1; can1-100; GAL+; Nab2-CBP-TEV-protA::TRP1 | This study |
| *NAB2-TAP Δhpr1* | MATa; ura3-1; trp1-1; his3-11,15; leu2-3,112; ade2-1; can1-100; GAL+; Nab2-CBP-TEV-protA::TRP1; hpr1::HIS3 | This study |
| *osTIR HPR1-AID NAB2-TAP* | URA3:::osTIR Hpr1-AID-6HA (plasmid 2352)::Hyg; Nab2-CBP-TEV-protA::TRP1 | This study |
| *MEX67-TAP* | MATa; ura3-1; trp1-1; his3-11,15; leu2-3,112; ade2-1; can1-100; GAL+; Mex67-CBP-TEV-protA::TRP | This study |
| *MEX67-TAP Δhpr1* | MATa; ura3-1; trp1-1; his3-11,15; leu2-3,112; ade2-1; can1-100; GAL+; Tho2-9myc::KanMX4; hpr1::HIS3; Mex67-CBP-TEV-protA::TRP | This study |
| *YRA1-TAP* | MATa; ura3-1; trp1-1; his3-11,15; leu2-3,112; ade2-1; can1-100; GAL+; Yra1-CBP-TEV-protA::TRP | This study |
| *YRA1-TAP Δhpr1* | MATa; ura3-1; trp1-1; his3-11,15; leu2-3,112; ade2-1; can1-100; GAL+; Tho2-9myc::KanMX4; hpr1::HIS3; Yra1-CBP-TEV-protA::TRP | This study |
| *THO2-9myc TEX1-6HA MFT1-3myc HPR1-6HA* | MATa; ura3-1; trp1-1; his3-11,15; leu2-3,112; ade2-1; can1-100; GAL+; Tho2-9myc::KanMX4; Tex1-6HA::Leu; Mft1-3myc::Trp; Hpr1-6HA::His | This study |
| *THO2-9myc TEX1-6HA MFT1-3myc HPR1-6HA THP2-TAP* | MATa; ura3-1; trp1-1; his3-11,15; leu2-3,112; ade2-1; can1-100; GAL+; Tho2-9myc::KanMX4; Tex1-6HA::Leu; Mft1-3myc::Trp; Hpr1-6HA::His; Thp2-TAP::Ura | This study |
| *THO2-9myc TEX1-6HA MFT1-3myc HPR1-6HA THP2-TAP Δhpr1* | MATa; ura3-1; trp1-1; his3-11,15; leu2-3,112; ade2-1; can1-100; GAL+; Tho2-9myc::KanMX4; hpr1::HIS3; Tex1-6HA::Leu; Mft1-3myc::Trp; Thp2-TAP::Ura | This study |
| *HPR1-AID THO2-9myc TEX1-6HA MFT1-9myc THP2-TAP* | URA3::osTIR; Hpr1-AID-6HA::Hyg; Tho2-9myc::KanMX4; Tex1-6HA::Leu; Mft1-9myc::His Thp2-Trp | This study |

**Supplementary Table 2. Plasmids**

| Name | Description | Reference |
|---|---|---|
| pBS1479 | For genomic C-terminal TAP-tagging (CBP-TEV-2x protein A), TRP1-KL | 4 |
| pYM14 | For genomic C-terminal 6xHA-tagging, KanMX4 | Euroscarf |
| pRS425 | High copy plasmid for overexpression, Leu marker | 5 |
| pRS425-*YRA1* | ORF + 500 bp of 5' and 500 bp of 3' UTR of *YRA1* was cloned into pRS425 | This study |
| pRS425-*MEX67* | ORF + 500 bp of 5' and 500 bp of 3' UTR of *MEX67* was cloned into pRS425 | This study |
| pRS425-*MEX67-6HA* | ORF of *MEX67* was C-terminal tagged with 6HA; + 500 bp of 5' and 500 bp of 3' UTR | This study |
| pRS425-*NAB2* | ORF + 500 bp of 5' and 500 bp of 3' UTR of *NAB2* was cloned into pRS425 | This study |
| pRS425-*NPL3* | ORF + 500 bp of 5' and 500 bp of 3' UTR of *NPL3* was cloned into pRS425 | 6 |

**Supplementary Table 3. Primers**

Primers used for cloning:

| Name | Sequence (5'→3') |
|---|---|
| OE NAB2 vector fwr | AAAATTATTTAATGGTTGATGAATTCCTGCAGCCCGG |
| OE NAB2 vector rev | TGTAATGAACCACACGATCGGATATCAAGCTTATCGATACCGTCG |
| OE NAB2 fragment fwr | GTATCGATAAGCTTGATATCCGATCGTGTGGTTCATTACA |
| OE NAB2 fragment rev | CCCCCGGGCTGCAGGAATTCATCAACCATTAAATAATTTTGTACACTTATAATAACAC |
| OE YRA1 vector fwr | TCATAATAGTTTTTTGAATTCCTGCAGCCCGGG |
| OE YRA1 vector rev | CGGGGAAATCCAATTGATATCAAGCTTATCGATACCGTCGAC |
| OE YRA1 fragment fwr | GATAAGCTTGATATCAATTGGATTTCCCCGAACAGC |
| OE YRA1 fragment rev | GGGCTGCAGGAATTCAAAAAACTATTATGATAACCTTGCTCAAGTCAAAC |
| OE MEX67 vector fwr | CTTGGACCGTAAAACTGCCTGCAGCCCGGGGGATCCACT |
| OE MEX67 vector rev | AAAATTTGGAGGACGAATTCGATATCAAGCTTATCGATACCGT |
| OE MEX67 fragment fwr | GCTTGATATCGAATTCGTCCTCCAAATTTTGCACCTC |
| OE MEX67 fragment rev | GATCCCCCGGGCTGCAGGCAGTTTTACGGTCCAAGCGG |
| MEX67-HA vector fwr | CTCGAATTCATCGATTAATGATATTGTTCCCTGTTTCAGCCG |
| MEX67-HA vector rev | GACCTGCAGCGTACGGAACTGCACAAATGCTTCTCTAGG |
| MEX67-HA fragment fwr | TAAACTGTATATTTTTTGTGATACTGTGCGGCTGAAACAGGGAACAATATCATTAATCGATGAATTCGAGCTCG |
| MEX67- HA fragment rev | AAAGGGTTTTCAGAGTAGCATGAATGGCATCCCTAGAGAAGCATTTGTGCAGTTCCGTACGCTGCAGGTCGAC |

Primers used for qPCRs after ChIP and RIP assays:

| Name | Sequence (5'→3') |
|---|---|
| NTR fwd[7] | TGCGTACAAAAAGTGTCAAGAGATT |
| NTR rev[7] | ATGCGCAAGAAGGTGCCTAT |

| | |
|---|---|
| *PMA1* 3’ fwd | CAGAGCTGCTGGTCCATTCTG |
| *PMA1* 3’ rev | GAAGACGGCACCAGCCAAT |
| *CCW12* 3’ fwd | TGAAGCTCCAAAGAACACCACC |
| *CCW12* 3’ rev | AGCAGCAGCACCAGTGTAAG |
| *YEF3* 3’ fwd | TCTGGTCACAACTGGGTTAGTG |
| *YEF3* 3’ rev | GCAATCTTGTTACCCATAGCATCGA |
| *ILV5* 3’ fwd | TGGTACCCAATCTTCAAGAATGC |
| *ILV5* 3’ rev | ACCGTTCTTGGTAGATTCGTACA |
| *RPL9B* 3’ fwd | AGGACGAAATCGTCTTATCTGGT |
| *RPL9B* 3’ rev | CAGATTTGTTGCAAGTCAGCGG |
| *DBP2* 3’ fwd | CTTCACCGAACAAAACAAAGGTT |
| *DBP2* 3’ rev | TCGGGAGGAATATTTTGATTAGCT |
| *ACT1* 3’ fwd | ATCATGAAGTGTGATGTCGATGTC |
| *ACT1* 3’ rev | ATGGTGGTACCACCGGACATAA |
| *RPS14B* 3’ fwd | AAGACCCCAGGACCAGGTG |
| *RPS14B* 3’ rev | GATACGGCCAATCCTCAAACCAG |
| *PRL28* 3’ fwd | TGGAAGCCAGTCTTGAACTTGG |
| *PRL28* 3’ rev | TTGGTCTCTCTTGTCTTCTGGGA |
